# Loss of vacuolar acidity results in iron sulfur cluster defects and divergent homeostatic responses during aging in *Saccharomyces cerevisiae*

**DOI:** 10.1101/2020.01.05.895433

**Authors:** Kenneth L. Chen, Toby N. Ven, Matthew M. Crane, Matthew L.C. Brunner, Adrian K. Pun, Kathleen L. Helget, Katherine Brower, Dexter E. Chen, Ha Doan, Justin D. Dillard-Telm, Ellen Huynh, Yen-Chi Feng, Zili Yan, Alexandra Golubeva, Roy A. Hsu, Raheem Knight, Jessie Levin, Vesal Mobasher, Michael Muir, Victor Omokehinde, Corey Screws, Esin Tunali, Rachael K. Tran, Luz Valdez, Edward Yang, Scott R. Kennedy, Alan J. Herr, Matt Kaeberlein, Brian M. Wasko

**Affiliations:** Department of Pathology, University of Washington, Seattle, WA, USA 98195; Department of Biology and Biotechnology, University of Houston-Clear Lake, Houston, TX USA 77058

## Abstract

The loss of vacuolar/lysosomal acidity is an early event during aging that has been linked to mitochondrial dysfunction. However, it is unclear how loss of vacuolar acidity results in age-related dysfunction. Through unbiased genetic screens, we determined that increased iron uptake can suppress the mitochondrial respiratory deficiency phenotype of yeast *vma* mutants, which have lost vacuolar acidity due to genetic disruption of the vacuolar ATPase proton pump. Yeast *vma* mutants exhibited nuclear localization of Aft1, which turns on the iron regulon in response to iron sulfur cluster (ISC) deficiency. This led us to find that loss of vacuolar acidity with age in wildtype yeast causes ISC defects and a DNA damage response. Using microfluidics to investigate aging at the single cell level, we observe grossly divergent trajectories of iron homeostasis within an isogenic and environmentally homogeneous population. One subpopulation of cells fails to mount the expected compensatory iron regulon gene expression program, and suffers progressively severe ISC deficiency with little to no activation of the iron regulon. In contrast, other cells show robust iron regulon activity with limited ISC deficiency, which allows extended passage and survival through a period of genomic instability during aging. These divergent trajectories suggest that iron regulation and ISC homeostasis represent a possible target for aging interventions.

## INTRODUCTION

Many studies of biological aging are performed with only a few age-points. While these types of studies have revealed a multitude of biological processes that become dysfunctional with age, they are unable to reveal the sequence, kinetics, and penetrance of age-associated changes. Defining these parameters is necessary in order to better understand the network of failures that results in age-associated pathological physiology. In the budding yeast (*Saccharomyces cerevisiae*), most aging studies have also been performed on populations of cells, lacking the resolution to identify differences within individual cells. Recent developments in microfluidic device technology have allowed for the characterization of yeast replicative aging with whole-lifespan breadth and single-cell resolution (Chen et al., 2017; Crane and Kaeberlein, 2018; Chen et al., 2019). This can allow for the identification of early drivers of aging as well as an understanding of the kinetics and penetrance of these changes, and information about their correlations with ultimate lifespan at single-cell granularity. This information may yield richer insight needed for the development of early interventions to combat age-associated diseases.

The lysosome/vacuole is a central node of cellular metabolism, with critical roles in nutrient sensing and storage, metal homeostasis, organelle maintenance, and protein degradation (Li and Kane, 2009). Accordingly, lysosomal dysfunction underlies many clinically significant genetic and degenerative diseases (Carmona-Gutierrez et al., 2016; Perera and Zoncu, 2016). The lysosome/vacuole is maintained at an acidic pH, which is necessary for proper organelle function. Recent work has shown that the loss of lysosomal/vacuolar acidity is an evolutionarily conserved early life driver of aging and mitochondrial dysfunction (Baxi et al., 2017; Ghavidel et al., 2018; Hughes and Gottschling, 2012), and interventions that increase vacuolar acidity have been found to extend lifespan in yeast (Sasikumar et al., 2019).

The acidic environment of the lysosomal/vacuolar lumen is generated by a highly conserved multi-subunit proton pump, the vacuolar ATPase (V-ATPase). In mammals, genetic ablation of the V-ATPase function is lethal (Inoue et al., 1999). However, in the budding yeast *Saccharomyces cerevisiae*, deletion of vacuolar ATPase subunits or dedicated assembly factors (*vma* mutants) is tolerated but results in a variety of physiological dysfunctions. In addition to alkalization of the vacuole, *vma* mutants exhibit respiratory incompetence (Eide et al., 1993); substantially reduced replicative lifespan (McCormick et al., 2015; Schleit et al., 2013); as well as sensitivity to elevated calcium levels (Ohya et al., 1986), alkaline pH (Serrano et al., 2004), iron depletion (Davis-Kaplan et al., 2004), reactive oxygen species (Milgrom et al., 2007), and DNA damage (Liao et al., 2007). How loss of V-ATPase activity and the resulting alterations in cellular pH homeostasis cause these pleiotropic phenotypes and limit lifespan remains unclear.

In both yeast and metazoans, lysosomal activity is necessary for the maintenance of iron homeostasis (Kidane et al., 2006; Milgrom et al., 2007). Iron is essential for many cellular functions and exists in various forms in the cell, including diiron, heme, and iron-sulfur clusters (ISCs). In yeast, the iron regulon is a coordinated gene expression program that is activated by iron deficiency and results in increased iron uptake and redistributed iron usage (Philpott and Smith, 2013). Interestingly, this program is triggered not by low total iron levels in the cell, but by reduced iron sulfur cluster (ISC) availability (Chen et al., 2004). ISCs are ancient protein prosthetic groups used in a wealth of essential cellular processes including DNA replication and repair, amino acid production, protein translation, and the electron transport chain (Lill et al., 2012). ISC production is dependent on a series of mitochondrial iron-transfer steps (Lill et al., 2012), and it has been speculated that they are the ultimate reason for the persistence of mitochondria in anaerobic eukaryotes. Indeed, mutations in proteins that utilize or assemble iron sulfur clusters underlie a plethora of devastating genetic cancer syndromes, hematopoietic abnormalities, and neurological diseases (Lill et al., 2012). Additionally, during conditions of ISC deficiency, genome maintenance is impaired (Díaz de la Loza et al., 2011; Pijuan et al., 2015; Veatch et al., 2009), mimicking a critical evolutionarily conserved hallmark of aging (López-Otín et al., 2013).

We hypothesized that determining how vacuolar acidity results in the pleiotropic defects and respiratory incompetence of *vma* mutants would provide insight into how age-associated loss of vacuolar acidity impacts cellular function during aging. Our results indicate that disruption of iron homeostasis is an important hub tying perturbed pH homeostasis and mitochondrial function. Stemming from this observation, we find that wildtype yeast display ISC defects during aging. Interestingly, we observe a divergence in iron-related homeostatic trajectories that cells can undergo during aging. A large subpopulation of cells appears unable to respond to the ISC-related crisis, while other cells mount the expected compensatory gene expression program that allows these cells to survive genomic instability and achieve a full lifespan potential.

## METHODS

### Reagents

Yeast strains used in this study are haploid and derived from the BY4741/4742 (S288C derived) background and genotypes are shown in Table S1. YEP media consisted of 2% Bacto Yeast Extract (Difco), 1% Bacto Peptone (Difco), and 2% glucose (YPD, fermentative conditions) or 3% glycerol (YPG, respiratory conditions).

### Yeast multicopy suppression screen

The multicopy suppression screen was performed using a YEP13 library (2µ, LEU2 vector) containing 5-20kb inserts of yeast genomic DNA (Nasmyth and Reed, 1980). The YEP13 pooled library was transformed into *vma21*Δ and *vma13*Δ *vma21*Δ double mutants and plated onto media lacking leucine to select for plasmid transformation. Approximately 10,000 colonies were then replica plated onto YPG media. For colonies that grew, the plasmid was isolated from yeast using a Qiagen plasmid prep kit (with glass beads added to lysis buffer) and the plasmid was transformed into *E. coli*, isolated (Qiagen) and sequenced with primers flanking the insertion site to identify the genomic region contained on the plasmid. For validation, *FET4* was cloned with ∼400bp of the upstream promoter into the pAG425 (resulting plasmid named pBMW182a) backbone using the Yeast Gateway System Vectors (obtained from Addgene) (Alberti et al., 2007).

### Yeast spontaneous and UV-mediated suppression screen

For the spontaneous genetic suppressor screen, *vma21*Δ cells were grown to log phase and then plated onto YPG plates. For the UV-mediated suppression screen, *vma21*Δ cells were exposed to UV radiation in an amount empirically determined to kill ∼50% of cells using a UV Stratalinker, and cells were plated on YPG media. Clones that were identified as suppressors were validated by rechecking growth on YPG. To identify the causative suppressing mutation, one strain was whole genome sequenced as described below, and subsequently other strains were Sanger sequenced at the *ROX1* locus.

### Whole genome sequencing

We used pooled linkage analysis to identify the mutation underlying the suppressor mutation (Birkeland et al., 2010). We first crossed the haploid mutant (BW1151) to a wildtype haploid. We then dissected tetrads from this strain and identified suppressor and non-suppressor spore clones based on their growth phenotypes on YPG. We grew overnight cultures from 10 of each class of segregant and counted the cells using a hemocytometer. We then combined 10^8^ cells from each culture into two pools corresponding to the two genotypes. We purified genomic DNA from the pooled cells using a Quick-DNA Fungal/Bacterial Miniprep kit (Zymo Research). We performed whole genome sequencing of the two pools essentially as described (Lee et al., 2019), except that we used a variant frequency cut-off of 0.8 to identify SNPs. There was a single SNP in the suppressor pool, which was present in 20 out of 20 reads and completely absent from the non-suppressed pool. The mutation maps to the *ROX1* locus on chromosome 16 and causes an A to T substitution in the first position of the fourth codon (AAA), creating a premature TAA stop codon (*rox1*-A10T).

### Yeast growth assays

Yeast growth assays on solid agar containing media were performed similarly to as previously described (Wasko et al., 2009). Briefly, yeast strains were transferred to into water and diluted to identical optical densities (OD600). Cells were aliquoted into 96 well microplates, five-fold serially diluted in water, and 4.5 µl of each sample were pipetted onto the indicated media type using a multi-channel pipette. After drying, plates were incubated at 30°C for 2-5 days.

### Aconitase activity assay

Aconitase activity assay was based on a previously described method (Tong and Rouault, 2006). Briefly, cell extracts were incubated with 0.2 mM phenazine methosulfate, 0.5 mM MTT (3-(4,5-Dimethylthiazol-2-yl)-2,5-Diphenyltetrazolium Bromide), 0.25 mM NADP, 2.5 mM cis-aconitic acid, and 0.4 U/ml isocitrate dehydrogenase and absorbance at 450 nm was quantified using a plate reader. An isocitrate standard was used to generate a standard curve and experimental data was collected with absorbance values that fell within the range of the linear standard curve. The BCA assay was used to determine equal protein amounts to add from each sample.

### Flow cytometry

Cells were grown to log phase in the indicated condition and cells were spun down, transferred into water, and stored on ice briefly until flow cytometry was performed with a FACSCanto II flow cytometer at the University of Washington Pathology Flow Cytometry Core Facility. Excitation was using a 488 nm laser and emission was monitored with a 502 long pass filter and a 530/30 filter. Three biological replicates were measured, each of which consisted of 20,000 cells per condition.

### Standard fluorescent microscopy

Wildtype or *vma21*Δ cells expressing Aft1-GFP were grown at 30°C to log phase in liquid YPD and then transferred to YPD in the presence or absence of 100 µg/ml bathophenanthrolinedisulfonic acid (BPS) or 1 mM iron ammonium sulfate for 2 hours at 30°C. Cells were then transferred to synthetic media (to reduce background fluorescence) with 2% glucose with or without 100 µg/ml BPS or 1 mM iron ammonium sulfate and imaged promptly after the media was changed.

### Replicative lifespan

Yeast replicative lifespan and statistical analysis were performed similar to as previously described (Wasko et al., 2013). Cells were patched to fresh YPD plates containing indicated compounds and grown overnight at 30°C. Virgin daughter cells of 20-40 cells per condition were selected for replicative lifespan microdissection analysis.

### Microfluidics and Fluorescence Microscopy

Cells were imaged using a PDMS microfluidic flow chamber with cell traps modified to increase mother cell retention and introduction of multiple chambers to isolate cells of different genotypes in identical environments. Cells were loaded according to previously published methods (Crane et al., 2014). A volumetric flow rate of 1-12 µL/min per chamber was used. Flow rate was initialized at a low rate to prevent ejection of smaller young mother cells from traps and increased throughout the experiment to reduce cell clogging and to maintain the trapping of large older mother cells. Cells were imaged using a Nikon Ti-2000 microscope with a 40X oil immersion objective, 1.3 NA using the Nikon Perfect Focus System. An enclosed incubation chamber was used to maintain a stable environment at 30 C. An LED illumination system (Excelitas 110-LED) was used to provide consistent excitation energies, with illumination triggered by camera shutter to prevent excess exposure. Images were acquired using a Hammamatsu Orca Flash 4.0 V2. The camera and stage were controlled by in-house software written in Matlab® and Micromanager. Images were corrected for illumination artifacts. To correct for single pixel biases, 1,000 images were acquired with no illumination, and individual pixel means were determined. To correct for flatness of field, fluorescent dye was added to a microfluidic device. 1,000 images were acquired each with a small offset in the x and y positions (to compensate for microfluidic trap features). Images were dilated, and the median value at each location was used. For every image, pixel-level bias was subtracted and values were multiplied by a flatness of field correction factor. Images were acquired at 5 min intervals for bright-field. Fluorescence imaging was acquired at 30 minute or 1 hr intervals. For bright-field imaging, 3 z-sections were 3.5 µm intervals. For the fluorescence, 3-7 z-sections were acquired. GFP images were acquired using a Chroma ET49002 filter set. mRuby2 and mCherry images were acquired using a Chroma ET49306 filter set. For pHluorin2 ratiometric imaging, the Chroma GFP ET49002 filter set and a custom ET405/40X excitation and ET525/50m emission filter set (Chroma) were used. Following data acquisition, cells were segmented and tracked using previously published software (Bakker et al., 2018). Divisions were scored by eye, and errors in cell segmentation and tracking were corrected manually. For Fit2 mean fluorescence, values of maximum projection were interpreted to reflect total protein levels. For vacuolar acidity, for each cell the brightness of the two fluorescence channels were equalized and summed, and the brightest 5% of pixels were used. The mean of the pixel-level ratio of these channels was used as a measure of vacuolar acidity. For Rad52 foci presence, the value of the brightest 9-pixel square divided by the brightest 2.5% of the cell was used. A threshold was determined by visual inspection of foci-containing and foci-free cells (1.1 for GFP, 1.05 for mCherry). For Rps2 nuclear retention, the value of the brightest 9 pixel square divided by the mean brightness of the cell was used.

Cells were inoculated into SC media (Sunrise Biosciences) with 2% dextrose and grown overnight (∼12-24 hours) until log phase. BSA was added immediately prior to loading to prevent adherence to PDMS. During experiments, SC media with 2% dextrose was used, and cells were imaged for 60-80 hrs.

## RESULTS

### Iron rescues pleiotropic phenotypes following disrupted V-ATPase function

Previous studies suggested that the loss of vacuolar acidity limits lifespan through mitochondrial dysfunction (Hughes and Gottschling, 2012). In order to identify links between vacuolar acidity and mitochondrial function, we performed a multi-copy screen for suppressors of the mitochondrial respiratory deficiency phenotype in *vma* mutants. We found that overexpression of the *FET4* gene encoding for a low-affinity iron transporter allowed *vma21*Δ cells to grow under respiratory conditions (Figure 1a). An additional screen for spontaneous and UV-mediated suppression of the *vma21*Δ respiratory deficiency yielded numerous loss-of-function alleles in the *ROX1* gene (Figure 1b and Table S2). Another genetic screen for suppressors of *vma21*Δ sensitivity to an iron chelator also yielded loss of function *ROX1* alleles (Table S3). Rox1p is a transcriptional repressor of genes involved in the yeast hypoxic response, so loss of *ROX1* results in activation of yeast hypoxic responsive genes. *FET4 is* one of the most upregulated targets in *rox1Δ* mutants (Jensen and Culotta, 2002; Ter Linde and Steensma, 2002; Waters and Eide, 2002), and *FET4* was required for suppression of *vma21*Δ respiratory deficiency by mutation of *ROX1* (Figure 1c).

**Figure 1.**
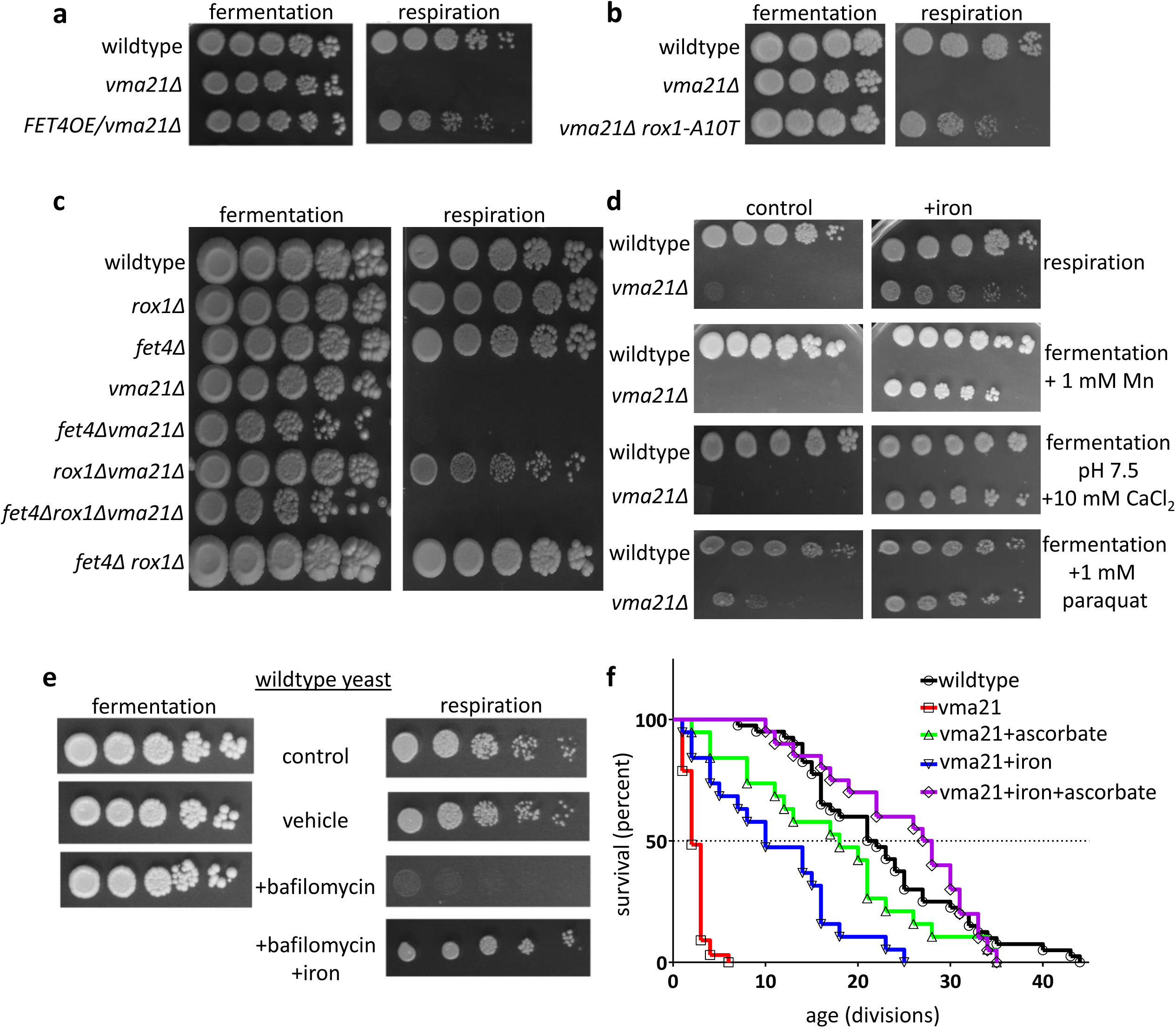
Enhanced iron uptake or additional iron allows respiratory growth following impairment of the V-ATPase. (**A**) Overexpression of *FET4* from a high copy (2µ) plasmid rescues growth of *vma21*Δ mutants under respiratory conditions. (**B**) A *vma21*Δ mutant strain containing a suppressing loss of function mutation in the *ROX1* gene (rox1-A10T) rescues the *vma21*Δ respiratory deficiency phenotype. (**C**) *FET4* is required for *rox1Δ* rescue of *vma21Δ* respiratory growth. (**D**) Supplemental iron (500 µM ferrous ammonium sulfate) rescues pleiotropic phenotypes of *vma21*Δ mutants. (**E**) Chemical inhibition of the V-ATPase with 3 µM bafilomycin A1 preferentially impairs growth under respiratory conditions and iron rescues this phenotype. (**F**) Replicative lifespan under fermentative growth conditions (YPD media) of wild type or *vma21*Δ mutants with or without 20 mM sodium ascorbate and/or 0.5 mM iron II sulfate. Where indicated, fermentative growth is on YPD media containing 2% glucose, and non-fermentative respiratory growth conditions is growth on YPG media containing 3% glycerol. Lifespan statistics are shown in Table S7.

Since Fet4 is an iron importer, we also tested supplementation of the growth media with iron, which also resulted in suppression of the *vma21*Δ respiratory deficiency (Figure 1d). This iron-mediated rescue of *vma* respiratory growth occurred with different forms of iron (II or III) and in multiple *vma* mutants lacking either cytosolic (V_1_) or membrane associated (V_0_) subunits of the V-ATPase (Figure S1a). Interestingly, iron supplementation additionally rescued other pleiotropic phenotypes of *vma21*Δ mutants including sensitivity to: elevated pH and calcium, manganese, and oxidative stress induced by paraquat (Figure 1d). Iron supplementation also rescued respiratory defects in wildtype yeast when the V-ATPase is chemically inhibited by bafilomycin (Figure 1e), suggesting that both acute and chronic inhibition of the V-ATPase impairs mitochondrial function by altering iron homeostasis. The short replicative lifespan of *vma21*Δ and *vma13*Δ mutants was additionally rescued by addition of iron and/or sodium ascorbate (Figure 1f and Figure S1b). Sodium ascorbate is an antioxidant that can reduce iron from the ferric (Fe^3+^) to ferrous (Fe^2+^) form (de Silva et al., 1997). The addition of sodium ascorbate diminished the iron regulon activation in *vma21*Δ mutants (Figure S1c), rescued the sensitivity of *vma21*Δ mutants to an iron chelator, and appeared to qualitatively increase the growth rate of wildtype yeast in the presence of an iron chelator (Figure S1d). This ability of sodium ascorbate to impact iron homeostasis may occur in three ways. Conceivably, ascorbate may have beneficial effects by reducing iron in the media (making it more soluble and bioavailable), by reducing iron intracellularly, and/or by acting directly as an intracellular antioxidant.

### Reduced V-ATPase function results in iron dyshomeostasis

In yeast, the response to low intracellular iron levels is mediated by nuclear localization of the transcription factor Aft1, which activates the iron regulon, transcription of a suite of genes involved in iron assimilation (e.g., *FET3*, *FTR1*, and *FIT2*). Using an Aft1 responsive transcriptional reporter expressing GFP under the control of the *FIT2* promoter (Diab and Kane, 2013), *vma21*Δ mutants displayed increased levels of GFP fluorescence compared to wild type, which was reduced by the addition of iron (Figure 2a). There is also nuclear localization of Aft1-GFP in *vma21Δ* cells, and exogenous iron reduced the nuclear localization of GFP tagged Aft1 in *vma21Δ* cells (Figure 2b). Although *vma21*Δ cells clearly induce the iron starvation response, we failed to detect any significant change in total intracellular iron levels (Figure 2c), which is consistent with prior reports for other *vma* mutants (Diab and Kane, 2013; Szczypka et al., 1997).

**Figure 2.**
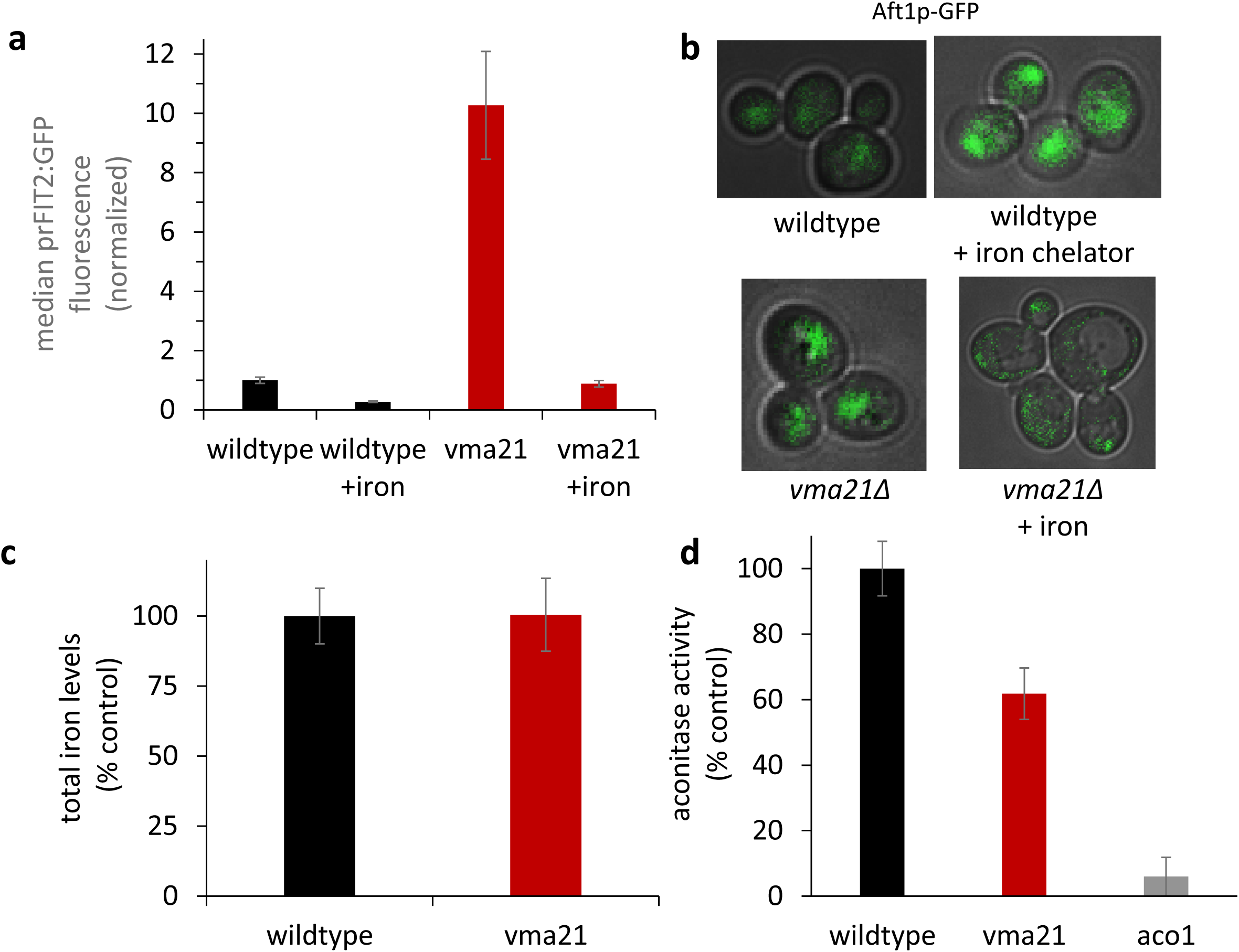
Iron dyshomeostasis occurs following disruption of the V-ATPase. (**A**) Flow cytometry analysis of GFP levels under control of the Aft1 responsive FIT2 promoter of wild type and *vma21*Δ mutants grown under fermentative conditions (YPD media) with or without 1 mM iron. Values are the mean of 3 samples, each consisting of 20,000 cells per condition. Statistics are shown in Table S8. Error bars represent the standard deviation of the 3 sample means. (**B**) Fluorescent microscopy images of wild type or *vma21*Δ mutants in the presence or absence of 100 µg/ml Bathophenanthrolinedisulfonic acid (BPS, iron chelator) or 1 mM iron ammonium sulfate. (**C**) Iron levels were measured in wildtype and *vma21*Δ cells grown in YPD media. No statistical difference was found using Student’s t-test. n=3, error bars represent standard deviation. (**D**) Aconitase activity was measured for wildtype cells, *vma21*Δ, and *aco1*Δ mutants. Each group is statistically significantly different from each other group by ANOVA and Bonferroni’s multiple comparison test P<0.001. n=3, error bars represent standard deviation.

Nuclear localization of Aft1 and activation of the iron regulon are triggered by deficits in iron sulfur cluster availability (Rutherford et al., 2005), suggesting the possibility that ISCs become limiting in *vma* cells. Aconitase is a mitochondrial ISC containing enzyme in the TCA cycle, and aconitase activity was reduced in *vma21*Δ cells (Figure 2d). consistent with a prior observation in *vma2* mutants (Diab and Kane, 2013).

Taken together, these observations suggested the possibility that disruption of iron homeostasis, and specifically, a deficiency in ISC availability, may fundamentally underlie some of the pleiotropic defects of *vma* mutant cells. Moreover, these observations suggest that since aging is accompanied by loss of vacuolar acidity, alterations in iron homeostasis and ISC status may occur during aging.

### Aging is characterized by a heterogeneous loss of vacuolar acidity coupled to an incompletely penetrant compensatory iron homeostatic response

To characterize the loss of vacuolar acidity during replicative aging, we tagged the vacuolar-localized carboxypeptidase Prc1 (Huh et al., 2003) with a ratiometric pH-sensitive fluorescent protein pHluorin2 (Mahon, 2011). Imaging these cells in a microfluidic device with live-cell fluorescence imaging indicated that loss of vacuolar acidity begins essentially at the start of life (Figure 3a). By comparing different genetic strains, previous studies have indicated that loss of vacuolar acidity limits lifespan (Ghavidel et al., 2018; Hughes and Gottschling, 2012). To test whether vacuolar acidity was a risk factor for early death in a wildtype isogenic population, we monitored changes in vacuolar acidity during early life (ages 0-12 divisions). In concordance with previous observations, the cells which experienced a faster rate of vacuolar acidity loss also exhibited a shorter lifespan (Figure 3b, Table S4), indicating that loss of vacuolar acidity is a predictor of replicative lifespan for individual cells within an isogenic population.

**Figure 3.**
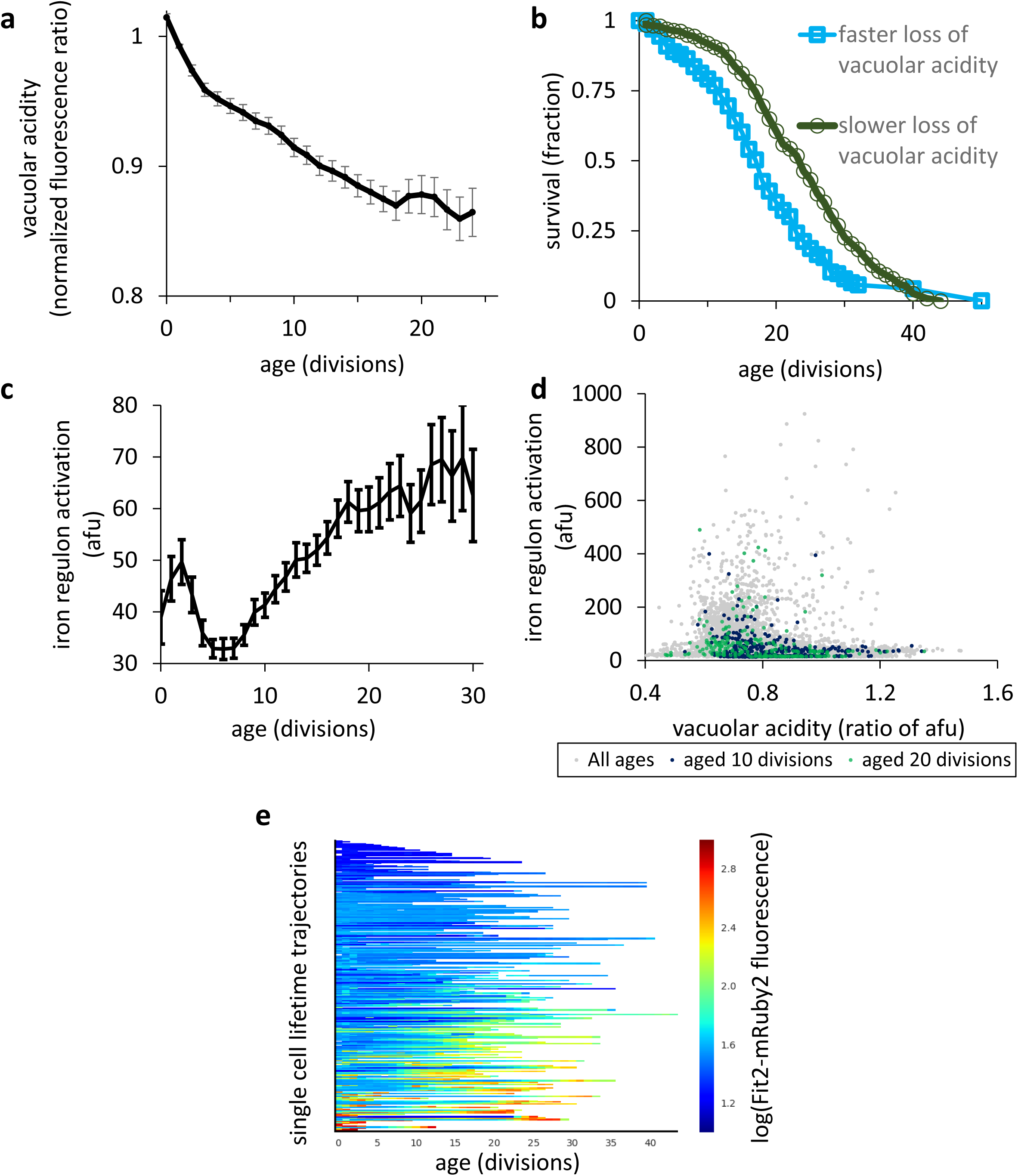
Age-associated decline in vacuolar acidity is predictive of replicative lifespan and is associated with iron regulon activity which increases with age. However, a large subset of cells displays little to no iron regulon activity during aging. (**A**) Vacuolar acidity trend during aging, as measured by fluorescence ratio of vacuole-localized Prc1-pHluorin2. Acidity is normalized per cell to young cell value (average before the second division). Pearson r = −0.259, p = 1 x 10^-149^, n = 9857 cell-divisions, error bars are standard error of the mean (SEM). (**B**) Survival curves for the population partitioned into two groups by the rate of vacuolar acidity loss during early life (0-12 divisions). The rate of vacuolar acidity loss rate for each cell is calculated using the slope of the least-squares regression line through the acidity values during divisions 0-12. Population is split in half using the median rate of vacuolar acidity loss. Fast rate of decline in vacuolar acidity is associated with shorter lifespan, logrank p = 2.22 x 10^-16^, n = 289 cells per group. (**C**) Age associated trend of population average iron regulon activity (Fit2-mRuby2 fluorescence). Iron regulon activity increases with age, Pearson r = 0.13, p < 10^-38^. (D) catter plot of iron regulon activity (Fit2-mRuby2 fluorescence) and vacuolar acidity for all cells at all ages. Iron regulon activity is correlated with lower vacuolar acidity even when controlled for age (All ages: Spearman ρ = −0.21, p < 10^-4^; age 10: Spearman ρ = −0.17, p = 0.0017; age 20: Spearman ρ = −0.38, p < 10^-4^). (**E**) Single cell lifetime trajectories of measured iron regulon activity. Color values are log(Fit2-mRuby2 fluorescence). Many cells show limited to no iron regulon activation.

Our genetic results suggested that aged cells may activate the iron regulon due to deficiency in ISCs. To test this hypothesis, we created a strain with C-terminal fluorescent protein tags for both vacuolar acidity (Prc1-pHluorin2) and an iron regulon reporter gene (Diab and Kane, 2013) (Fit2-mRuby2). Averaged across the population, we observed a progressive increase in iron regulon activity during replicative aging (Figure 3c). On a single-cell level, iron regulon activation was also associated with decreased vacuolar acidity even when controlled for age (Figure 3d, Table S5). However, many aged cells with reduced vacuolar acidity showed little to no activation of the iron regulon (Figure 3d). Indeed, many cells failed to activate the iron regulon to any appreciable degree during aging (Figure 3e). Thus, population-level iron regulon activation during aging was driven by cells that induced Fit2 several orders of magnitude above cells that failed to activate the iron regulon.

### Divergent trajectories emerge during aging: active iron regulon/limited ISC deficiency and inactive iron regulon/runaway ISC deficiency

Upon observing that the age-associated iron regulon activation was only partially penetrant, we sought to evaluate the single-cell levels of ISC deficiency during aging. Is the activation of a strong iron regulon response during aging protective against ISC deficiency, or would it indicate a more severe deficiency in ISCs? To answer this question, we measured the activity of the essential ISC protein Rli1 during aging. Rli1 transports the small ribosomal subunit protein Rps2 out of the nucleus into the cytoplasm, and Rli1 activity has been used as a reporter of overall ISC sufficiency (Kispal et al., 2005). Under conditions of ISC deficiency, Rps2 accumulates in the nucleus, so that the fraction of Rps2 localized to the nucleus is a measure of the insufficiency of active Rli1 and of ISCs generally (Kispal et al., 2005). We created a reporter strain with both an iron regulon reporter (Fit2-mRuby2) and a GFP-tagged Rps2 expressed under the GPD promoter on chromosome I. In this strain, ISC insufficiency results in the appearance of a bright fluorescent dot representing Rps2-GFP sequestered in the nucleus (Kispal et al., 2005). By observing this strain during aging in a microfluidic device, we found an overall increase in the level of Rps2-GFP foci with age, suggesting ISC insufficiency during aging (Figure 4a). Unexpectedly, we observed that in middle-aged and old cells (>12 divisions), severity of ISC deficiency was inversely correlated to the level of iron regulon activation (Figure 4b). We assigned each cell into one of two groups—iron regulon active or iron regulon inactive—based on its maximum observed fluorescence level from Fit2-mRuby2 during aging. When we measured the overall progression of ISC deficiency during aging in the iron regulon active and inactive subpopulations, we found the ISC deficiency increased similarly for both groups until roughly division 12. As cells entered middle age, the iron regulon-active cells plateaued in ISC deficiency, while iron regulon-inactive cells became progressively more ISC deficient (Figure 4c). Visualizing the paths of all individual cells until death reveals that iron regulon activation and ISC deficiency define two clearly divergent iron homeostasis trajectories during aging (Figures 4d, 4e): iron regulon-activation/ISC-sufficiency and iron regulon-inactivity/ISC-deficiency.

**Figure 4.**
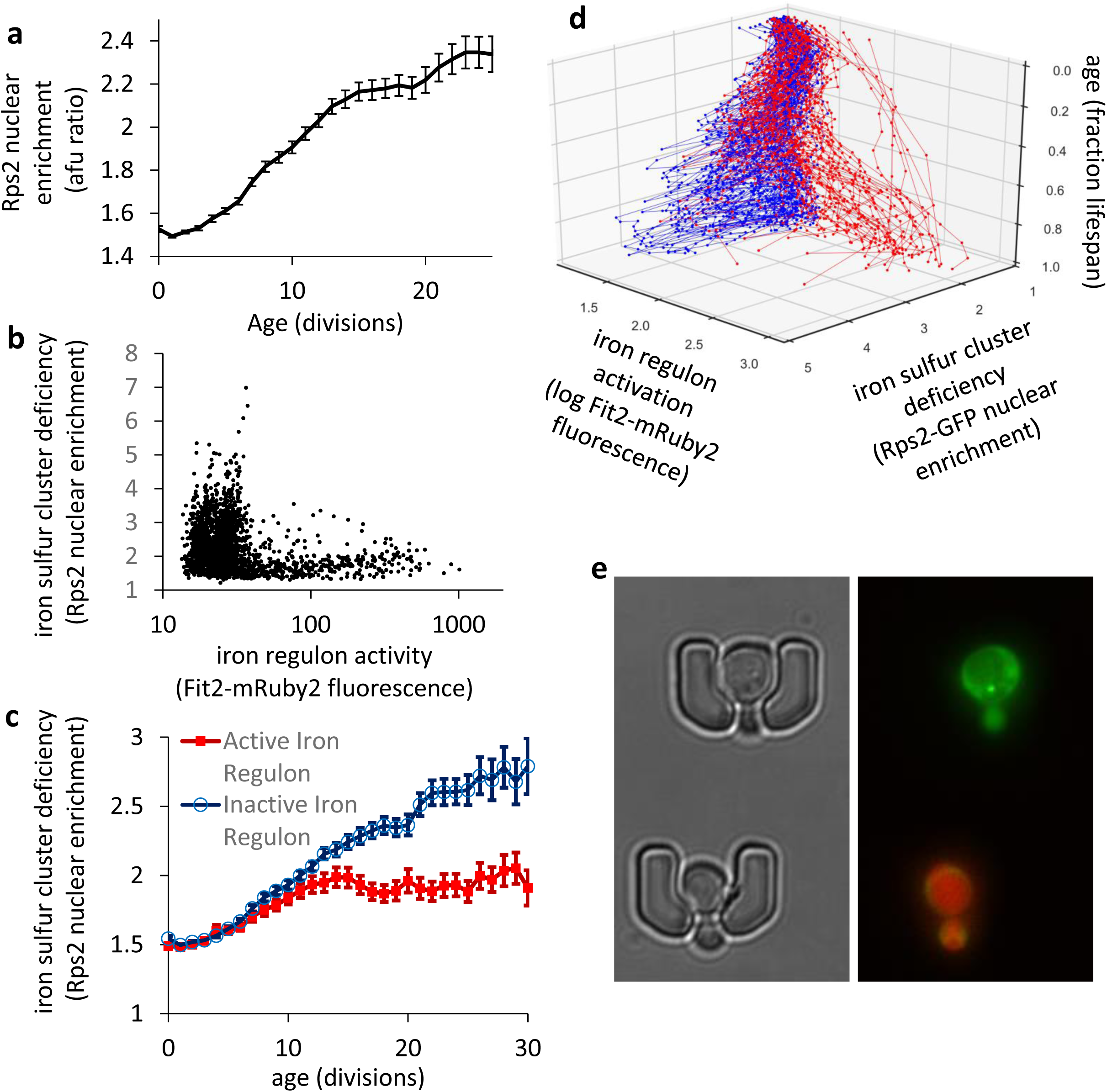
Iron sulfur clusters become deficient during aging. Activation of iron regulon is inversely correlated with iron sulfur cluster deficiency. Cells with robust iron regulon activity during aging generally follow a trajectory of limited iron sulfur cluster deficiency. n=209 total cells. (**A**) Aging trend of population mean Iron sulfur cluster insufficiency (Rps2 nuclear enrichment). Iron sulfur cluster insufficiency increases during aging. Pearson r = 0.52, p < 10^-4^. Error bars are SEM. (**B**) Scatter plot of iron sulfur cluster insufficiency (nuclear enrichment of Rps2-GFP) and iron regulon activation (Fit2-mRuby2 fluorescence) in aged cells (all cells and ages > 12 divisions). Spearman ρ = −0.26, p < 10^-4^, n=189 (cells that lived 12 generations or more). (**C**) Aging trend of Iron sulfur cluster deficiency for iron regulon competent and incompetent cells during aging. Iron regulon active cells are defined as having a maximum Fit2 fluorescence level that is 3x the median baseline. Iron regulon inactive cells are not more iron-starved during early life, but have less severe iron sulfur cluster deficiency from middle to old age (p < 0.05 for ages 15-30). n_active iron regulon_= 59, n_inactive iron regulon_= 150. Error bars are SEM. (**D**) Single cell trajectories of iron sulfur cluster deficiency and iron regulon activity during aging. Cells display divergent iron metabolism trajectories during aging: Red: iron regulon active with limited iron sulfur cluster deficiency. Blue: no iron regulon activation. n=209. (**E**) Example aged cells in adjacent traps from each iron metabolism trajectory. Top cell in trap: Iron sulfur cluster deficiency (Rps2-GFP nuclear retention visible as bright green dot) with little iron regulon activation. Bottom cell in trap: Robust iron regulon activation (Fit2-mRuby2) with no visible Rps2-GFP nuclear retention.

### Loss of vacuolar acidity during aging is associated with elevated indications of genomic instability

Since vacuolar acidity is important for ISC sufficiency and ISC proteins mediate DNA replication and repair, the loss of vacuolar acidity may be related to genomic instability during aging. To investigate this hypothesis, we studied the development of genome instability during replicative aging in a microfluidic device using a strain with our Prc1-pH2 vacuolar acidity reporter and an mCherry-tagged Rad52, a central mediator of the DNA damage response (DDR). Upon DNA double-strand breaks, Rad52 directs the formation of a gigadalton-sized protein complex (usually one per haploid genome), which functions as processing centers for DNA repair through homologous recombination (Gasior et al., 1998). When Rad52 is fluorescently tagged, these DNA repair centers are visible as distinct subcellular foci (Lisby et al., 2001). Thus, the population frequency of Rad52 foci has been commonly used as a reporter for DNA damage severity (Alvaro et al., 2007; Novarina et al., 2017).

We and other researchers (Novarina et al., 2017) observed a population-wide increase in the frequency of activation of Rad52-mediated DNA repair during aging (Figure 5a), indicating there may be a general increase in spontaneous DNA damage. Cells which experience spontaneous DNA damage earlier in life also had a shorter replicative lifespan (Table S6), suggesting that DNA damage may be one physiological factor that limits life. We also observed an age-controlled correlation between lower vacuolar acidity and DNA damage during aging. When looking at both activation of the DNA damage response pathway and vacuolar acidity, we observed that during middle to old-age, cells in which the DNA damage response was activated had reduced vacuolar acidity compared to age-matched cells without a DNA damage response (Figure 5b). Furthermore, a faster initial drop in vacuolar acidity during early life was correlated with an earlier appearance of DNA damage during aging (Figure 5c). This is consistent with the idea that the age-associated loss of vacuolar acidity is a possible upstream driver of genome instability during aging.

**Figure 5.**
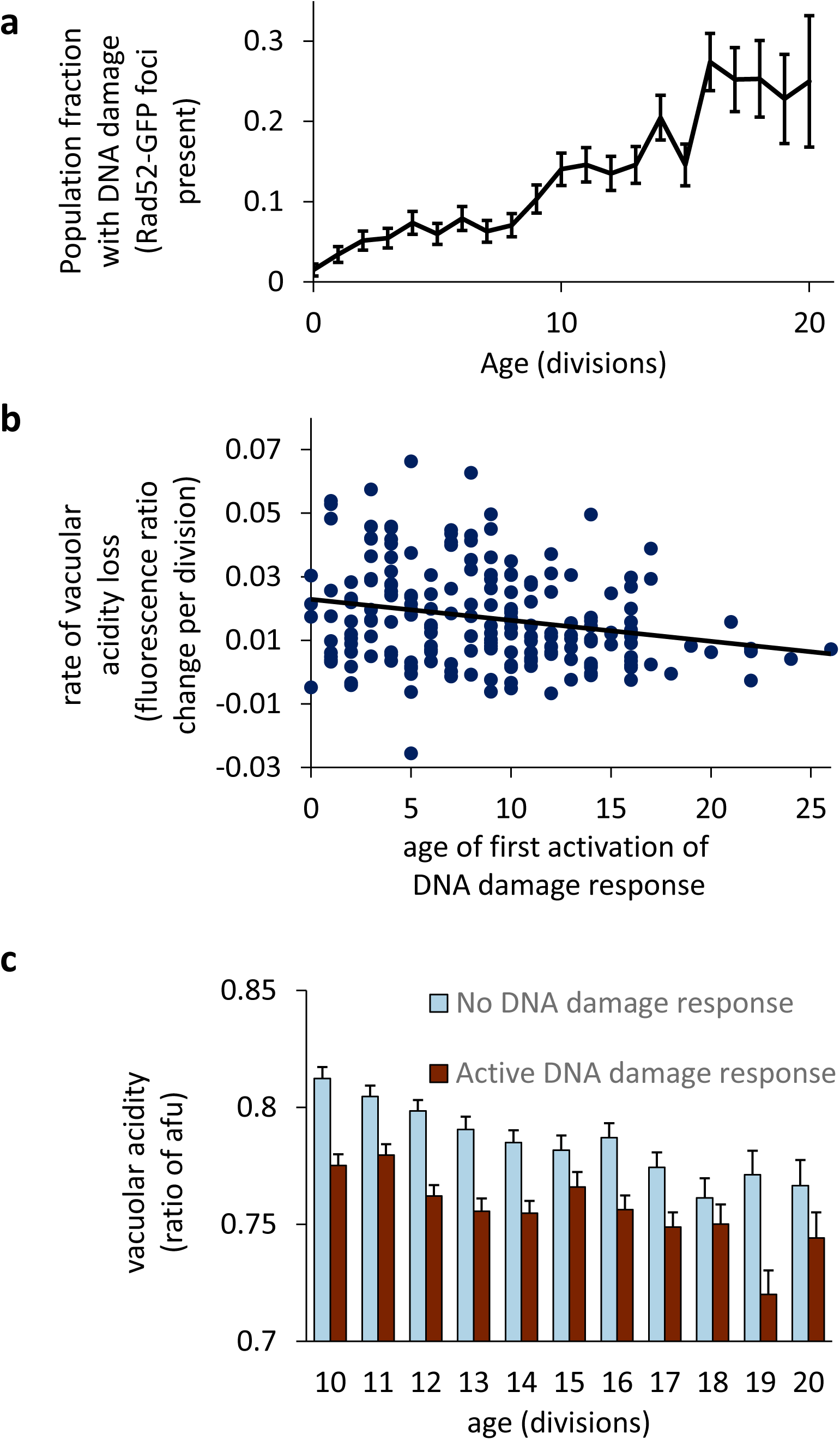
Activation of the DNA damage response increases during aging, is correlated with the loss of vacuolar acidity, and is predictive of a shorter replicative lifespan. (**A**) Aging trend of DNA damage response activation as measured by presence of Rad52-GFP foci. As cells age the fraction of cells that activate the DNA damage response increases, Pearson r = 0.27, p < 10^-4^, n=266 cells. Error bars are SEM. (**B**) Scatter plot of age of first DNA damage response activation (Rad52-GFP foci) and rate of vacuolar acidity loss in early life for cells which activate the DNA damage response at any point during life (ages 0-12 divisions). n=244 cells that developed foci. Faster loss of vacuolar acidity during early life (ages 1-12 divisions) correlates with earlier appearance of DNA damage response. Pearson r = −0.23, p = 0.0006. (**C**) Comparison of vacuolar acidity for cells with and without activated DNA damage response (Rad52-GFP foci) at various ages. n=266. Presence of activated DNA damage response (Rad52-GFP foci) is associated with lower vacuolar acidity when controlled for age. All comparisons ranksum p< 0.05 except for ages 15: p = 0.089, 18: p = 0.45, and 20: p = 0.20. Error bars are SEM.

### The iron homeostasis trajectory of a cell is associated with its genome stability

Iron metabolism and genome maintenance are interconnected, as ISCs are necessary for both DNA replication and many types of DNA repair (Netz et al., 2014). Accordingly, impairment of ISC production has been shown to cause genome instability (Veatch et al., 2009) and reduced survival in genotoxic conditions (Pijuan et al., 2015). Because of this, differences in genome stability between the two subpopulations of divergent iron metabolism aging trajectories were evaluated. Strains were generated with both an iron regulon reporter (Fit2-mRuby2) and a DNA damage reporter (Rad52-GFP) and replicative aging was measured in a microfluidic device. Cells were separated into two groups, either iron regulon-active or iron regulon-inactive based on the maximum iron regulon activity observed during aging. Cells which activated the iron regulon during aging survived for longer after the initial observation of spontaneous DNA damage (Figure 6a and 6b). Iron regulon active cells also underwent more cell divisions during which the DNA damage response had been activated (Figure 6c). We speculate that because the iron regulon-active cells experienced limited ISC deficiency during aging, these cells may be better able to survive spontaneous age-related DNA damage by marshalling the appropriate ISC-dependent DNA repair processes.

**Figure 6.**
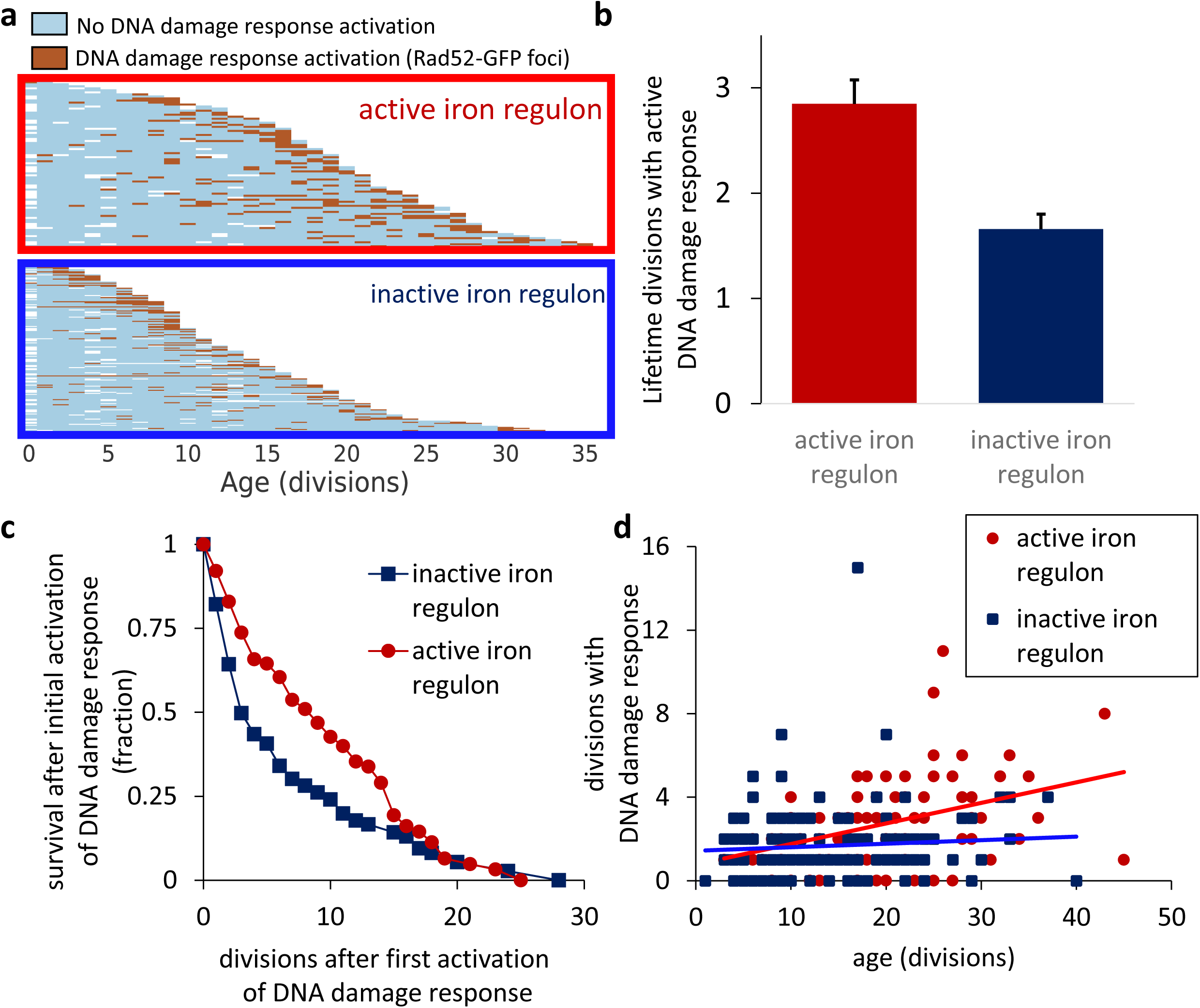
Iron regulon active cells survive longer after the first activation of the DNA damage response during aging and undergo more lifetime divisions during which the DNA damage response (Rad52-GFP foci formation) is activated. Total lifespan of iron regulon active cells is highly correlated to the number of lifetime divisions during which the DNA damage response is activated. Within the iron regulon inactive group, longer-lived cells do not undergo any more divisions with DNA damage than shorter lived cells. This suggests that within the iron regulon competent subpopulation, the longest-lived cells are those with the most robust ability to survive age-associated DNA damage. Within the iron regulon incompetent subpopulation, the longest-lived cells are those with minimal age-associated DNA damage. (**A**) Single cell trajectories of DNA damage response activation during aging. Top: Cells that activate the iron regulon during aging (maximum lifetime Fit2-mCherry fluorescence > 3x median baseline level) Bottom: Cells that fail to activate the iron regulon during aging. Empty spaces indicate that data was not collected for a single cell at that age, e.g. due to temporary cell segmentation or tracking error or initial age of capture after first division. (**B**) Survival curves comparing remaining lifespan after first observed activation of the DNA damage response (Rad52-GFP foci formation) for iron regulon competent and incompetent cells. Cells that activate the iron regulon during aging have increased survival after the first activation of the DNA damage response logrank p = 0.018, n_active iron regulon_= 76, n_inactive iron regulon_= 129. Error bars are SEM. (**C**) Comparison of DNA damage response activation between iron regulon active and iron regulon inactive cells. During aging, cells that activate the iron regulon undergo more divisions during which the DNA damage response is activated (student’s t p < 10^-4^). (**D**) Scatter plots showing correlation between total lifespan and number of divisions during which the DNA damage response is activated. Longer lifespan of the iron regulon active cells is well-correlated (Pearson r = 0.41) with a higher number of divisions with an activated DNA damage response. Longer-lifespan in iron regulon inactive cells is not correlated (Pearson r = 0.08) with more divisions with DNA damage. Correlation coefficients are significantly different by Fisher’s Z-transform p = 0.01.

Interestingly, differing relationships between DNA damage response activation and longevity between these two iron metabolism sub-populations were observed. Within the iron regulon-active group, longer lifespan was correlated with additional divisions during which the DNA damage response was activated (Figure 6d). Within the iron regulon-inactive group, there was no correlation between longer lifespan and activation of the DNA damage response (Figure 6d). One explanation of these results is that the longest-lived cells within these groups may have different molecular processes underlying their survival. Within the iron regulon-active group, longevity may be determined by the robustness of the repair and survival responses to spontaneous DNA damage. Within the iron regulon-inactive group, with more severe ISC deficiency and potentially less capability to repair DNA damage, longevity may be more dependent on lower rates of spontaneous DNA damage during aging.

## DISCUSSION

Using a combination of traditional yeast genetics, microfluidics, and fluorescence microscopy, we have further characterized the cellular consequences of loss of vacuolar acidity, an early-life physiological change during aging. We found that loss of vacuolar acidity results in iron-dependent pleiotropic phenotypes of *vma* mutant cells. This suggests that iron dyshomeostasis is an important consequence following disruption of the V-ATPase. In *vma* mutants, there is both a loss of vacuolar acidity as well as cytosolic acidification (Martinez-Munoz and Kane, 2008). Intracellular acidification can damage iron-sulfur clusters (Follmann et al., 2009; Li et al., 2011), and cytosolic acidification by acetic acid treatment is sufficient to activate the iron regulon (Diab and Kane, 2013). This indicates that cytosolic acidification following loss of the V-ATPase may contribute to activating the iron regulon by damaging iron sulfur clusters. With age, pH near the cytoplasmic membrane has been reported to increase (Henderson et al., 2014), while cytosolic pH has been reported to acidify (Knieß and Mayer, 2016). It is possible that during aging, cytosolic acidification may result in damage to iron sulfur clusters. Yeast V-ATPase mutants display elevated levels of intracellular reactive oxygen species (ROS) (Milgrom et al., 2007) and ISCs are highly sensitive to degradation by ROS (Jang and Imlay, 2007). Interestingly, disruption of ISC production is itself also a source of ROS (Gomez et al., 2014), suggesting the possibility of vicious cycle during aging.

What specifically happens to iron homeostasis following alterations in cellular pH homeostasis remains to be determined. Yeast V-ATPase mutants exhibit defects in the maturation of Fet3, a high affinity iron uptake protein (Davis-Kaplan et al., 2004). However, *fet3* mutants do not fully phenocopy *vma* mutants, and we found no difference in total cellular iron levels in *vma21* cells, suggesting additional factors beyond reduced iron uptake lead to the pleiotropic phenotypes.

In rodent cultured hepatocytes, inhibition of the V-ATPase results in the release of chelatable iron from lysosomes, which is then taken up by mitochondria (Uchiyama et al., 2008). In aging mice, iron has also been found to accumulate in mitochondria (Seo et al., 2008). Disruption of mitochondrial iron-sulfur cluster biosynthesis in yeast results in the accumulation of iron within the mitochondria and activation of the iron regulon in spite of normal cytosolic iron levels (Chen et al., 2004). The iron regulon activation and iron levels we and others observe in *vma* mutants are consistent with these observations. However, since we observe that additional iron rescues a mitochondrial defect in *vma* mutants, a model of mitochondrial iron accumulation would seem to require that the mitochondria would harbor iron in a form that is not bioavailable. Future studies could assess the intracellular localization, trafficking, and redox state of iron in *vma* mutants in order to identify if iron is mislocalized and/or if the redox state of iron is aberrant, and these results could then generate hypotheses for what may occur following age-associated pH dyshomeostasis.

We found that wildtype aged yeast cells display a deficiency in iron sulfur clusters and an increased DNA damage response. Furthermore, both of these physiological declines correlate with vacuolar acidity loss when controlled for age. Unexpectedly, we found that only a subset of yeast cells can activate the expected iron regulon program in response to iron sulfur cluster deficiency during aging. Thus, an isogenic, environmentally homogeneous population of yeast cells follows divergent iron metabolism trajectories during aging. Iron regulon-active cells mount a robust activation of the iron regulon, resulting in limited ISC deficiency throughout life. In contrast, more than half of the population is unable to mount any meaningful iron regulon activity, resulting in a progressive worsening of ISC deficiency during aging. This divergence has implications on other measures of physiology during aging, including cellular responses to age-associated spontaneous DNA damage. We find that in general, iron regulon active cells are better able to survive and continue dividing during episodes of apparent DNA damage, with a strong correlation between longer lifespan and additional DNA damage episodes survived. In contrast, iron regulon-inactive cells appear sensitive to DNA damage, and longer-lived cells in this population appear to be those which generally escaped DNA damage early in life, requiring minimal activation of the DNA damage response. We have not attempted to fully characterize the differences between the iron regulon-active and iron regulon-inactive subpopulations, including the mechanistic details of DNA damage survival during aging. While the fidelity of DNA repair was not investigated, we observed an intriguing divergence during aging of single cells into a DNA damage-resistant and -vulnerable states underpinned by the activation of (or lack thereof) the iron regulon gene expression program.

The concept of cell-state divergence during aging that we describe has recently been characterized in terms of yeast mother and daughter cell morphology (Jin et al., 2019). Although no gene expression or mechanistic observations were made in that study, our shared observations suggest that the aging process may have characteristics reminiscent of Waddington’s landscape of development, where cells diverge into distinct aged states. The mechanistic underpinnings of these apparently stochastic paths of aging present a novel and intriguing area of study. We expect that aging trajectories may be more fully characterized by single-cell observation of additional gene expression or physiological parameters. While this study and Jin et al. (2019) made observations of different parameters, it seems likely that integration of these observations as well as those of other physiological variables will result in the identification of more divergent physiological variables during aging. Ultimately, whether the total multidimensional landscape of aging can be best understood as a finite array of valleys, or whether this analogy is better suited to a separate understanding of single parameters of aging is yet to be determined.

The loss of vacuolar acidity is an evolutionarily conserved aging phenotype identified thus far in yeast (Ghavidel et al., 2018; Hughes and Gottschling, 2012) and worms (Baxi et al., 2017). Future studies will need to directly assess whether loss of lysosomal acidity occurs in mammalian cell types with age or age-associated diseases, which could be testable using ratiometric pH sensors. If loss of lysosomal acidity occurs during mammalian aging, further studies could seek to determine if it is a driving force for age-associated iron accumulation, mitochondrial dysfunction, and/or ISC dysfunction. Impairment of lysosomal acidity has been tied to iron dyshomeostasis in mammalian cell lines (Lim et al., 2007; Miles et al., 2017; Schneider et al., 2015; Straud et al., 2010). ISC proteins function in a wide variety of physiological contexts and are highly conserved evolutionarily (Netz et al., 2014). Moreover, defects in ISC production or in specific ISC proteins underlie numerous genetic syndromes with progeroid characteristics as well as many age-associated diseases (Perera and Zoncu, 2016). We propose that a pH-dependent impairment of iron sulfur clusters with age may be a fundamental and evolutionarily conserved driver of clinically significant age-associated pathology.

