## Supplemental Figures and Tables for "Loss of vacuolar acidity results in iron sulfur cluster defects and divergent homeostatic responses during aging in *Saccharomyces cerevisiae*"

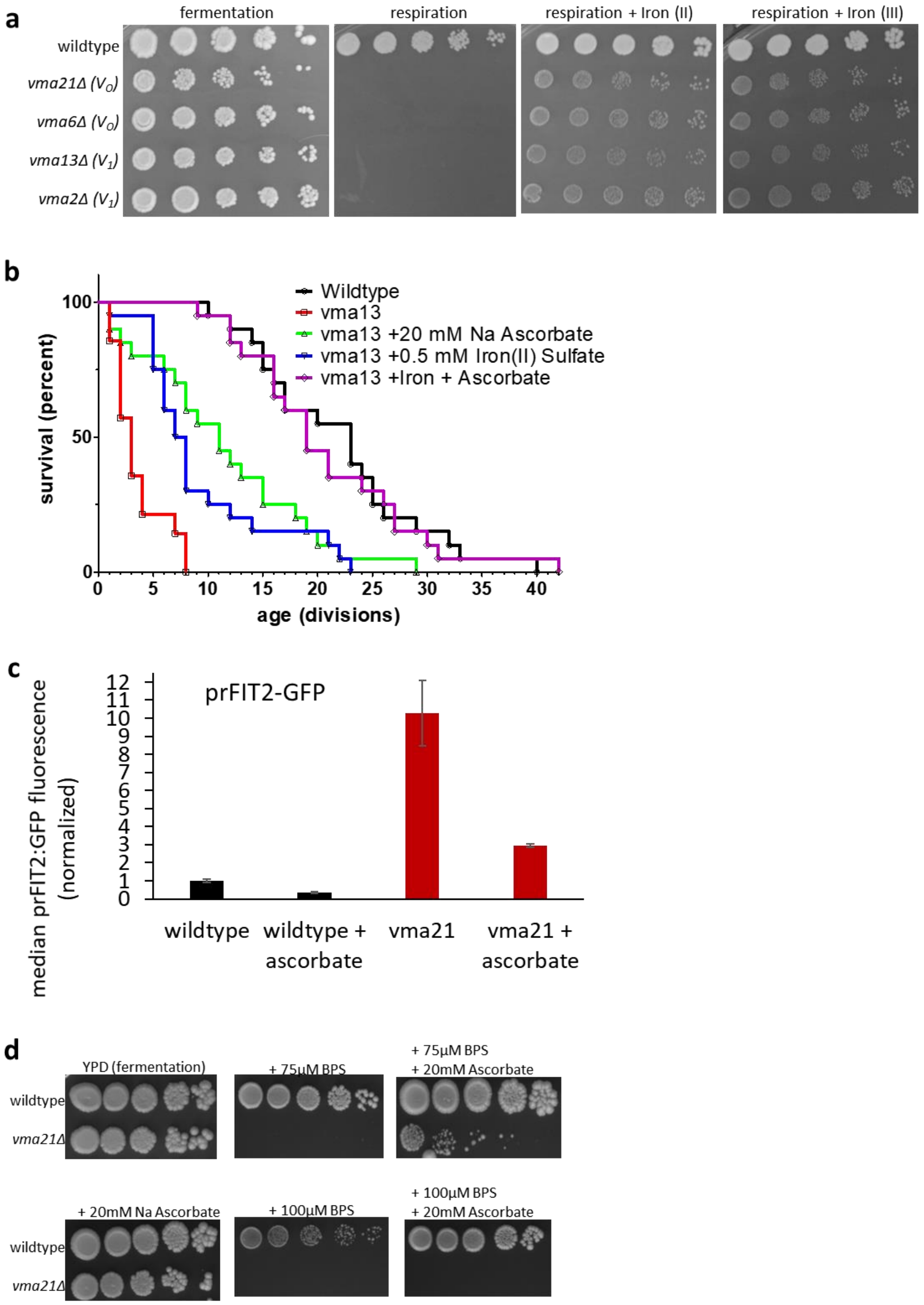


**Figure S1. Exogenous iron II or iron III rescues vma $V_0$ and $V_1$ mutants, and sodium ascorbate impacts iron homeostasis.** (A) Addition of 2.5 mM iron II sulfate or 2.5 mM iron III chloride rescues the growth defect of *vma21*, *vma6*, *vma13*, and *vma2* cells under respiratory conditions (YPG). (B) The shortened replicative lifespan of vma13 mutants is increased by iron and/or sodium ascorbate. (C) Fluorescence levels of GFP under control of the FIT2 promoter were measured by flow cytometry in wild-type yeast and *vma21*Δ mutants in the presence or absence 20 mM sodium ascorbate. n=3, with 20,000 cells per condition. (D) Sodium ascorbate rescues the sensitivity of *vma21*Δ mutants to an iron chelator, BPS.

| Strain | Genotype | Reference/Source |
|---|---|---|
| BY4741 | Wildtype, MATa ura3Δ his3Δ1 leu2Δ met15Δ | ThermoFisher Scientific |
| BY4742 | Wildtype, MATα ura3Δ his3Δ1 leu2Δ lys2Δ | ThermoFisher Scientific |
| BW1016 | MATα vma21::URA3 his3Δ1 leu2Δ lys2Δ | this study |
| BW1017 | MATa vma21::URA3 his3Δ1 leu2Δ met15Δ | this study |
| BW1008 | MATa FIT2pr::GFP | Diab and Kane. 2013. |
| BW1025 | MATa FIT2pr::GFP vma21::URA3 | this study |
| BW1134 | MATa vma21::URA3 vma13::KanMX | this study |
| BW1151 | MATa rox1-A10T | this study |
| BW1200 | MATa fet4::KanMX | ThermoFisher Scientific |
| BW1212 | MATa rox1::KanMX | this study |
| BW1218 | MATa vma21::URA3 rox1::KanMX | this study |
| BW1241 | MATa vma21::URA3 fet4::KanMX | this study |
| BW1245 | MATa vma21::URA3 fet4::KanMX rox1::KanMX | this study |
| BW1247 | MATa rox1::KanMX fet4::KanMX | this study |
| BW1322 | MATa AFT1-GFP:HIS3 | ThermoFisher Scientific |
| BW1325 | MATa vma21::URA3 AFT1-GFP:HIS3 | this study |
| aco1 | aco1::KanMX | ThermoFisher Scientific |
| KC385 | MATα prc1::PRC1-pHluorin2, HphNT1 fit2::FIT2-mRuby2, CaHIS5 | this study |
| KC620 | MATα pGPD::RPS2-GFP, URA3 fit2::FIT2-mRuby2, CaHIS5 | this study |
| KC631 | MATa prc1::PRC1-pHluorin2, HphNT1 rad52::RAD52-mCh, KanMX | this study |
| KC614 | MATa fit2::FIT2-mCherry, KanMX rad52::RAD52-GFP, HIS3 | this study |

**Table S1.** Yeast strains used in this study.

| Selection | # of strains | *ROX1* mutation | mutation type |
|---|---|---|---|
| Spontaneous, YPG | 2 | A10T | nonsense |
| Spontaneous, YPG | 3 | C346T | nonsense |
| Spontaneous, YPG | 1 | T1105G | nonstop |
| UV, YPG | 3 | A271T | nonsense |

**Table S2. Genetic suppressors of *vma* respiratory deficiency.**

| Selection | # of strains | *ROX1* mutation | mutation type |
|---|---|---|---|
| Spontaneous, BPS | 1 | G193T | nonsense |
| Spontaneous, BPS | 1 | T408 or T409 deletion | frameshift |

**Table S3. Genetic suppressors of *vma* BPS sensitivity.**

| Age | Pearson r | p value |
|---|---|---|
| 4 | 0.10 | 3.66E-04 |
| 5 | 0.11 | 1.47E-04 |
| 6 | 0.16 | 1.81E-07 |
| 7 | 0.17 | 3.46E-08 |
| 8 | 0.22 | 5.92E-13 |
| 9 | 0.24 | 4.01E-15 |
| 10 | 0.23 | 1.86E-13 |
| 11 | 0.26 | 1.57E-16 |
| 12 | 0.27 | 2.96E-16 |
| 13 | 0.21 | 1.10E-10 |
| 14 | 0.25 | 4.19E-13 |
| 15 | 0.23 | 7.90E-11 |
| 16 | 0.22 | 1.33E-09 |
| 17 | 0.23 | 4.84E-09 |
| 18 | 0.22 | 4.66E-08 |
| 19 | 0.21 | 8.24E-07 |
| 20 | 0.23 | 6.89E-07 |
| 21 | 0.22 | 7.28E-06 |
| 22 | 0.16 | 1.50E-03 |
| 23 | 0.19 | 3.02E-04 |
| 24 | 0.20 | 6.27E-04 |
| 25 | 0.16 | 1.20E-02 |
| 26 | 0.25 | 2.01E-04 |
| 27 | 0.22 | 2.10E-03 |
| 28 | 0.18 | 2.24E-02 |
| 29 | 0.23 | 8.43E-03 |
| 30 | 0.31 | 1.80E-03 |

**Table S4. Higher vacuolar acidity as measured by fluorescence ratio of Prc1-pHluorin2 is significantly correlated to higher remaining lifespan across the aging timeline.** Each row shows the correlation coefficient and statistical significance between remaining lifespan and vacuolar acidity at a particular age for all cells.

| Age | Spearman rho | p value |
|---|---|---|
| 10 | -0.16 | 1.49E-03 |
| 11 | -0.19 | 2.20E-04 |
| 12 | -0.22 | 4.38E-05 |
| 13 | -0.24 | 6.42E-06 |
| 14 | -0.29 | 7.90E-08 |
| 15 | -0.32 | 8.74E-09 |
| 16 | -0.39 | 1.31E-12 |
| 17 | -0.35 | 3.07E-10 |
| 18 | -0.35 | 3.04E-09 |
| 19 | -0.37 | 1.31E-09 |
| 20 | -0.38 | 4.86E-10 |
| 21 | -0.36 | 9.74E-09 |
| 22 | -0.36 | 6.27E-08 |
| 23 | -0.34 | 7.81E-07 |
| 24 | -0.35 | 2.28E-06 |
| 25 | -0.30 | 1.76E-04 |

**Table S5: Correlation between vacuolar acidity (Prc1-pH2 fluorescence ratio) and iron regulon activity (Fit2-mRuby2 fluorescence).** When controlled for age, vacuolar acidity and iron regulon activity are inversely correlated.

| Age | Hazard Ratio | p value |
|---|---|---|
| 1 | 1.688 | 0.091 |
| 2 | 1.387 | 0.173 |
| 3 | 1.513 | 0.044 |
| 4 | 1.591 | 0.009 |
| 5 | 1.575 | 0.006 |
| 6 | 1.591 | 0.003 |
| 7 | 1.571 | 0.003 |
| 8 | 1.558 | 0.003 |
| 9 | 1.540 | 0.002 |
| 10 | 1.364 | 0.025 |
| 11 | 1.335 | 0.034 |
| 12 | 1.278 | 0.069 |

**Table S6. Activation of the DNA damage response (measured by presence of Rad52-GFP foci) in early life is associated with shorter lifespan.** Table shows the cox proportional hazards ratio for cells with early activation of the DNA damage response vs cells with no early activation of the DNA damage response. Early activation is defined as activation of the DNA by the age shown in the first column.

| | Log-rank (Mantel-Cox) Test p value | Gehan-Breslow-Wilcoxon Test p value |
|---|---|---|
| wildtype vs vma21+iron+ascorbate | 0.6532 | 0.2637 |
| vma21 vs vma21+ascorbate | < 0.0001 | < 0.0001 |
| vma21 vs vma21+iron | < 0.0001 | < 0.0001 |
| vma21 vs vma21+iron+ascorbate | < 0.0001 | < 0.0001 |

**Table S7. Statistics for Figure 1f, determined using Prism GraphPad 5.**

| One-way analysis of variance | |
|---|---|
| P value | <0.0001 |
| P value summary | *** |

| Bonferroni's Multiple Comparison Test | Significant? P < 0.05? | Summary |
|---|---|---|
| Wildtype(wt) vs wt+iron | No | ns |
| wt vs vma21 | Yes | *** |
| wt vs vma21+iron | No | ns |
| wt+iron vs vma21 | Yes | *** |
| wt+iron vs vma21+iron | No | ns |
| vma21 vs vma21+iron | Yes | *** |

**Table S8. Statistics for Figure 2a, determined using Prism GraphPad 5.**
